# CodonRL: Multi-Objective Codon Sequence Optimization Using Demonstration-Guided Reinforcement Learning

**DOI:** 10.64898/2026.02.12.705465

**Authors:** Shiyi Du, Gün Kaynar, Jiayi Li, Zhaoyi You, Shijie Tang, Carl Kingsford

**Affiliations:** Carnegie Mellon University

## Abstract

Optimizing synonymous codon sequences to improve translation efficiency, RNA stability, and compositional properties is challenging because the search space grows exponentially with protein length and objectives interact through long range RNA structure. Dynamic programming-based methods can provide strong solutions for fixed objective combinations but are difficult to extend to additional constraints. Deep generative models require large-scale, high-quality mRNA sequence datasets for training, limiting applicability when such data are scarce. Reinforcement learning naturally handles sequential decision-making but faces challenges in codon optimization due to delayed rewards, large action spaces, and expensive structural evaluation. We present CodonRL, a reinforcement learning framework that learns a structural prior for mRNA design from efficient folding feedback and demonstration-guided replay, and then enables user-controlled multi-objective trade-offs during inference. CodonRL uses LinearFold for fast intermediate reward computation during training and ViennaRNA for final evaluation, warms up learning with expert sequences to accelerate convergence for global structure objectives, and introduces milestone-based intermediate rewards to address delayed feedback in long range optimization. On a benchmark of 55 human proteins, CodonRL outperforms GEMORNA, a state-of-the-art codon optimization method, across multiple metrics, achieving 9.5% higher codon adaptation index (CAI), 25.4 kcal/mol more favorable minimum free energy (MFE), and 3.4% lower uridine content on average, while improving codon stabilization coefficient (CSC) in over 90% of benchmark proteins under matched constraints. These gains translate into designs that are predicted to be more efficiently translated, more structurally stable, and less immunogenic, while supporting continuous objective reweighting at inference time.

## Introduction

The design of optimal codon sequences is a fundamental challenge in synthetic biology and biotechnology [Endy, 2005, Benner and Sismour, 2005]. While the genetic code’s degeneracy allows multiple codon sequences to encode the same protein, the choice of specific codons impacts protein expression levels, with variations spanning several orders of magnitude [Plotkin and Kudla, 2011, Perlak et al., 1991, Gustafsson et al., 2004]. This raises a fundamental question: how can we design codon sequences that simultaneously optimize expression, stability, and desired compositional features in a flexible and generalizable way? This optimization problem has critical implications for therapeutic protein production [Ward et al., 2011, Mauro, 2018], vaccine development [Zhang et al., 2023, Xia, 2021], and industrial biotechnology applications [Elena et al., 2014, Schmidt et al., 2023]. Recent mRNA vaccine efforts, including those developed for COVID-19, further highlight the practical importance of optimizing mRNA coding sequences under structural and compositional constraints [Chaudhary et al., 2021, Verbeke et al., 2022].

Traditional codon optimization approaches have focused on maximizing the codon adaptation index (CAI) to match host organism codon usage preferences [Sharp and Li, 1987]. Recent advances in RNA biology have revealed that mRNA secondary structure stability, quantified by minimum free energy (MFE), plays a role in determining expression levels [Presnyak et al., 2015, Hanson and Coller, 2018]. The inherent trade-off between these objectives (improving CAI often compromises MFE and vice versa) necessitates multi-objective optimization strategies [Şen et al., 2020, Chin et al., 2014].

Methods such as LinearDesign [Zhang et al., 2023] employ dynamic programming algorithms to navigate this optimization landscape. While effective, these approaches suffer from several limitations: (1) they are typically restricted to optimizing a limited set of metrics (MFE and CAI) and lack flexibility for incorporating additional design objectives or constraints; (2) they employ relatively fixed heuristic search strategies with limited capacity to explore diverse solutions; (3) they struggle with computational scalability for long protein sequences due to the *O*(*L*^3^) complexity of RNA folding algorithms (where *L* is the sequence length) [Zuker and Stiegler, 1981, McCaskill, 1990]. Deep generative models represent an alternative paradigm that has recently gained prominence. GEMORNA [Zhang et al., 2025], a transformer-based encoder-decoder architecture, demonstrates zero-shot generation capabilities and achieves substantial improvements in experimental validation. Nevertheless, this approach presents complementary constraints: model training requires large-scale, high-quality mRNA sequence datasets and considerable computational resources for pre-training; and the framework lacks explicit mechanisms for dynamic adjustment of multi-objective optimization trade-offs, limiting adaptability to diverse design specifications and constraints. Together with the limitations of dynamic programming approaches, this motivates methods that can learn strong structural preferences while still supporting flexible, user-directed multi-objective control.

Reinforcement learning (RL) has emerged as a powerful approach for sequential decision making in biological sequence design and has been successfully applied to problems ranging from DNA sequence optimization to protein engineering [Angermueller et al., 2019, Brookes et al., 2019]. The codon optimization problem exhibits a sequential and state-dependent structure, where each codon choice not only affects the immediate output but also influences structural stability, nucleotide composition, and translation efficiency. This delayed and interdependent reward landscape makes the problem particularly well-suited for reinforcement learning, which naturally handles long-range dependencies and credit assignment across decoding steps. However, applying RL to codon optimization presents unique challenges: sparse reward signals, high-dimensional action spaces, and the computational expense of evaluating RNA secondary structures at each step.

We present a novel reinforcement learning framework, called CodonRL, that addresses these challenges through several innovations (see also Figure 1):

**Figure 1:**
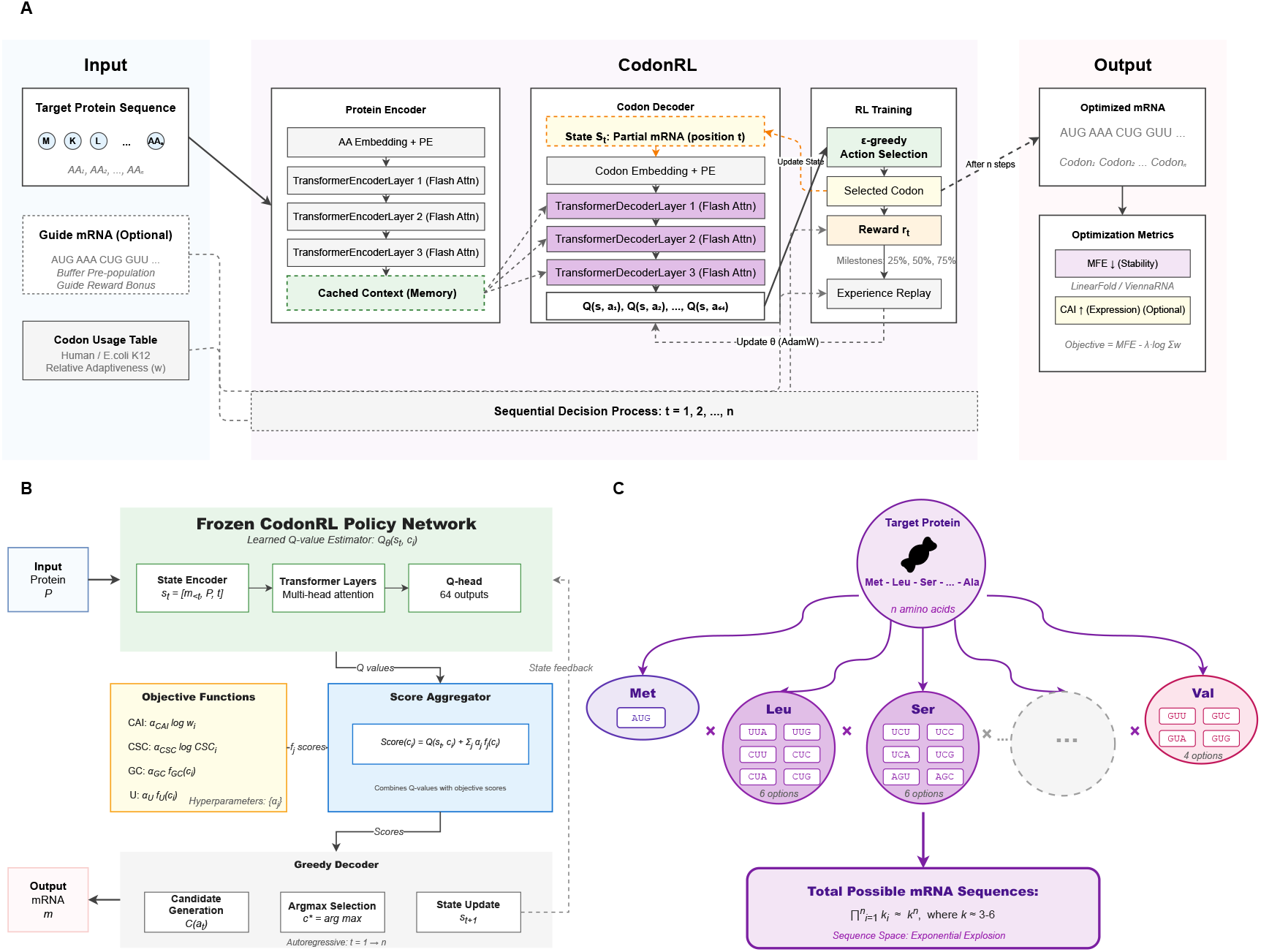
Overview of the CodonRL framework for mRNA sequence optimization. **(A)** Reinforcement learning-based codon optimization workflow. Left: Input protein sequence (AA_1_, AA_2_, …, AA_*n*_) with optional guide mRNA and species-specific codon usage table. Center: CodonRL architecture consists of a protein encoder (3-layer Transformer with flash attention) that generates cached context representations, and a codon decoder (3-layer Transformer) that autoregressively generates Q-values for all 64 possible codons at each position. The decoder receives partial mRNA state *s*_*t*_ and attends to protein context via cross-attention (dashed arrows). Right: RL training module performs *ϵ*-greedy action selection, computes rewards with milestone bonuses (25%, 50%, 75% completion), and updates the policy network *θ* via experience replay using AdamW optimizer. Sequential decision process iterates from *t* = 1 to *L* amino acids. Final sequences are evaluated on MFE (minimum free energy via LinearFold/ViennaRNA) and optional CAI metrics. **(B)** Inference-time decoding with frozen CodonRL policy. Top: Frozen policy network outputs Q-values Q_*θ*_(*s*_*t*_, *c*_*i*_) for each codon candidate. Middle left: Objective functions compute weighted scores for CAI (translation efficiency), CSC (codon stabilization coefficient), GC content, and U content, with tunable hyperparamete *{α*_*j*_ *}*. Middle right: Score aggregator combines Q-values with objective scores: Score(*c*_*i*_) = Q(*s*_*t*_, *c*_*i*_) + Σ*j α*_*j*_ *f*_*j*_ (*c*_*i*_). Bottom: Greedy decoder performs candidate generation, argmax selection, and autoregressive state updates to produce the final optimized mRNA sequence. **(C)** Combinatorial explosion of the mRNA sequence space. Each amino acid (e.g., Met, Leu, Ser, Val) can be encoded by multiple synonymous codons (1-6 options depending on degeneracy). The total number of possible mRNA sequences grows exponentially as 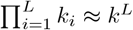 where *k ∈* [3, 6] represents the average codon degeneracy and *L* is the protein length, creating an astronomically large search space (e.g., ≥ 10^180^ sequences for a 300 amino acid protein) that necessitates advanced optimization strategies.

1. **Demonstration-guided learning:** We implement a pre-population mechanism that leverages high-quality mRNA sequences to initialize replay buffers and provide exploration bonuses with reinforcement learning exploration. Unlike typical codon optimization pipelines, we explicitly incorporate a small set of externally optimized sequences as expert demonstrations to warm-start learning and improve sample efficiency for global structural objectives.
2. **Efficient structural feedback:** We use LinearFold [Huang et al., 2019] for reward calculation during the training loop. By leveraging its *O*(*L*) complexity for intermediate feedback, we avoid the computational bottleneck of standard folding algorithms during optimization, reserving thermodynamic accuracy (e.g., ViennaRNA [Lorenz et al., 2011]) for final sequence evaluation.
3. **Reward densification mechanism:** We introduce milestone-based intermediate rewards at 25%, 50%, and 75% sequence completion to address the sparse reward problem inherent in long range optimization, providing dense feedback signals that guide learning throughout the generation process.
4. **Post-training multi-objective adaptation:** Unlike prevailing methods that optimize fixed metric combinations or produce single deterministic outputs, our RL framework enables dynamic control over design objectives through adaptive objective weighting at inference time.

We demonstrate CodonRL’s superior performance on a diverse set of proteins across multiple evaluation metrics including minimum free energy (MFE), codon adaptation index (CAI), codon stabilization coefficient (CSC), and uridine rate, outperforming GEMORNA on average with 9.5% higher CAI, 25.4 kcal/mol more favorable MFE, and 3.4% lower uridine content, while improving CSC in over 90% of benchmark proteins and providing greater design flexibility and interpretability. Source code for the method is available at https://github.com/Kingsford-Group/codonrl.

## Results

### Experimental Setup

#### Benchmark Dataset

We evaluate CodonRL on protein sequences (*n* = 55) from the LinearDesign UniProt benchmark [Zhang et al., 2023], all human sequences with *≤* 500 amino acids selected for computational tractability. The dataset spans diverse functional categories including receptors and signaling proteins, membrane transporters, enzymes, immune-related proteins, and proteins involved in RNA/protein synthesis, chromatin regulation, and cytoskeletal organization. This functional diversity ensures evaluation across diverse proteins with varying biochemical properties and cellular localizations.

#### Baseline Methods

We compare methods representing different optimization paradigms: (1) **Random sampling** from synonymous codons as a naive baseline; (2) **Natural**, the wild-type coding sequences;(3) **CAI-maximization**, a greedy algorithm that selects the highest-frequency codon at each position;(4)**LinearDesign** [Zhang et al., 2023] with *λ* = 0 (pure MFE optimization) and *λ* = 4 (balanced MFE-CAI trade-off); (5) **GEMORNA** [Zhang et al., 2025], a recent deep generative model for mRNA design; and (6) **CodonRL** (our method) with various tradeoffs between multiple objectives (*α* weights) and guidance from expert sequences. We construct the guidance using LinearDesign (*λ* = 0) to generate a series of sequences optimized in the MFE direction. This strategy bootstraps the RL framework with stable structures, as minimizing MFE (a global property) is usually harder for the RL framework to solve via cold-start exploration compared to other metrics (see Figure 6 for an ablation study demonstrating the performance gains from this guidance).

#### Evaluation Metrics

We assess sequence quality across five dimensions: (1) **Codon Adaptation Index (CAI)**, measuring translation efficiency through codon usage bias (higher is better); (2) **Minimum Free Energy (MFE)** in kcal/mol, quantifying mRNA secondary structure stability via ViennaRNA (more negative indicates greater stability). We verify the computational cost of accurate MFE calculation in Supplementary Figure 7, comparing ViennaRNA against faster alternatives like LinearFold; (3) **Codon Stabilization Coefficient (CSC)**, capturing codon pair-level stability preferences (higher is better) [Wu et al., 2019]; (4) **GC content (GC%)**, assessed for proximity to the optimal range; and (5) **Uridine content (U%)**, where lower values reduce innate immune activation for therapeutic applications. For multi-objective comparisons, we compute per-sequence improvements ΔMetric = Metric_CodonRL_ - Metric_baseline_ and report win rates (percentage of sequences where CodonRL outperforms the baseline).

### Global Design Space Comparison: CodonRL vs. GEMORNA

We compared CodonRL with GEMORNA to assess their relative ability to optimize multiple biologically meaningful mRNA properties. The analysis integrates sequence-level distributions, population-level geometric structure, and protein-wise paired comparisons (Figure 2).

**Figure 2:**
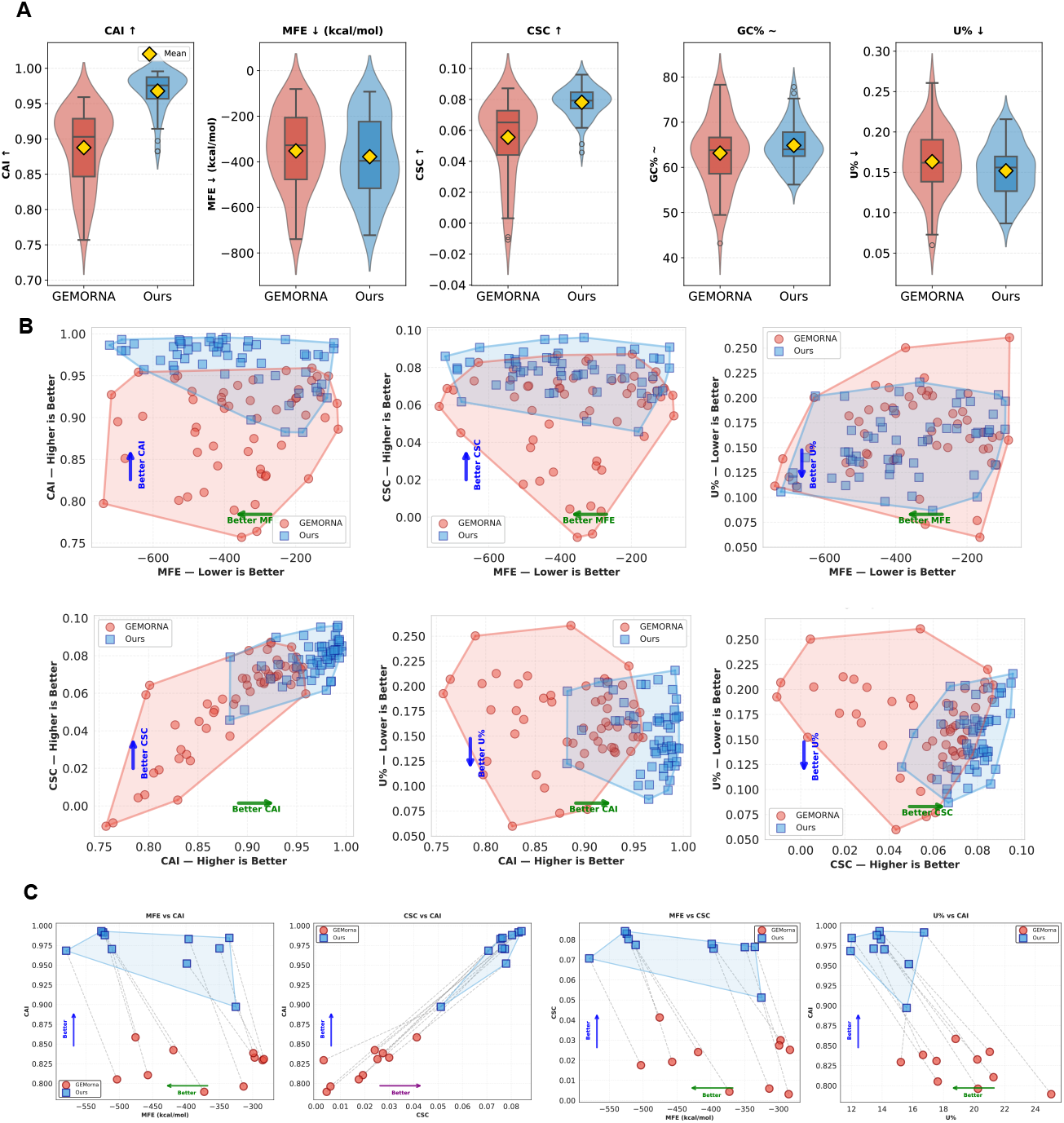
Global and local design-space comparison between GEMORNA and CodonRL. **(A)** Violin–box plots comparing the marginal distributions of five key sequence-level metrics between GEMORNA (red) and CodonRL (blue) across all benchmark proteins: codon adaptation index (CAI, higher is better), minimum free energy (MFE, kcal/mol; more negative indicates more stable secondary structure), codon-stability coefficient score (CSC, higher is better), GC content (GC%, maintained within a target range), and uridine fraction (U%, lower is preferred to reduce innate immune activation). For each metric, violins show that CodonRL systematically shifts the distributions toward higher CAI and CSC, more favorable (more negative) MFE, controlled GC%, and reduced U%. **(B)** Population-level 2D design spaces for all sequences, illustrating pairwise trade-offs between objectives. Each panel plots GEMORNA designs (red circles and convex hull) and CodonRL designs (blue squares and convex hull) for combinations of MFE with CAI, CSC, and U% (top row), and CAI with CSC and U% as well as CSC with U% (bottom row). Axis annotations and arrows indicate the “better” direction for each objective (e.g., MFE: left, CAI/CSC: up, U%: down). The expansion and shift of the blue convex hulls demonstrate that CodonRL enlarges and moves the attainable design space toward the multi-objective optimum, achieving sequences that are jointly more stable, more highly expressed, and compositionally better tuned than GEMORNA. **(C)** Per-sequence improvements for the top 10 proteins ranked by a composite multi-objective score. For each protein, paired GEMORNA (red circles) and CodonRL (blue squares) designs are shown in four 2D projections (MFE vs. CAI, CSC vs. CAI, MFE vs. CSC, and U% vs. CAI), with dashed gray line segments connecting matched designs and blue convex hulls summarizing the CodonRL front. The consistent movement of points along the annotated “better” directions demonstrates that, on a per-protein basis, CodonRL typically achieves simultaneous gains in structural stability, codon usage, and stability-related metrics while reducing U%, rather than merely redistributing trade-offs among objectives.

#### Systematic Shifts Across Biological Properties

Across the benchmark proteins, CodonRL yields higher CAI and CSC, more favorable (more negative) MFE, stable GC% centered near the target range, and modestly reduced U% relative to GEMORNA (Figure 2A). These improvements are broad rather than driven by a small subset of proteins: the interquartile ranges for CAI, MFE, and CSC shift substantially away from those of GEMORNA.

#### Expansion of the Attainable Design Space

CodonRL expands the attainable multi-objective design space relative to GEMORNA across pairwise trade-offs among CAI, MFE, CSC, and U% (Figure 2B). Across these projections, CodonRL reaches regions that GEMORNA does not, extending toward the Pareto-favorable direction for each objective pair. In contrast, GEMORNA concentrates designs within a narrower, correlated region of the landscape. This broader coverage indicates that CodonRL does not simply reweight existing trade-offs; it discovers sequence designs that jointly improve codon usage, structural stability, and composition in combinations that are largely absent from GEMORNA outputs.

#### Per-Protein Trajectory Analysis of Most-Improved Sequences

Figure 2C examines the ten proteins for which CodonRL achieves the largest multi-objective gains relative to GEMORNA. These cases are the proteins where GEMORNA’s designs fall furthest from the desired multi-objective optima and thus correspond to the sequences where improvement is most needed. Across all objective projections, the paired trajectories exhibit consistent increases in CAI and CSC and simultaneous decreases in MFE and U% from GEMORNA to CodonRL. This demonstrates that CodonRL does not derive its global advantages from small aggregated improvements across easy cases, but instead achieves substantial per-protein enhancements precisely in the sequences where GEMORNA performs poorly.

### Multi-Method Benchmarking and Objective Correlation Analysis

We situate CodonRL within a broad comparison of alternative sequence design strategies and show how different methods shape the correlation structure and local sequence properties across biological objectives (Figure 3).

**Figure 3:**
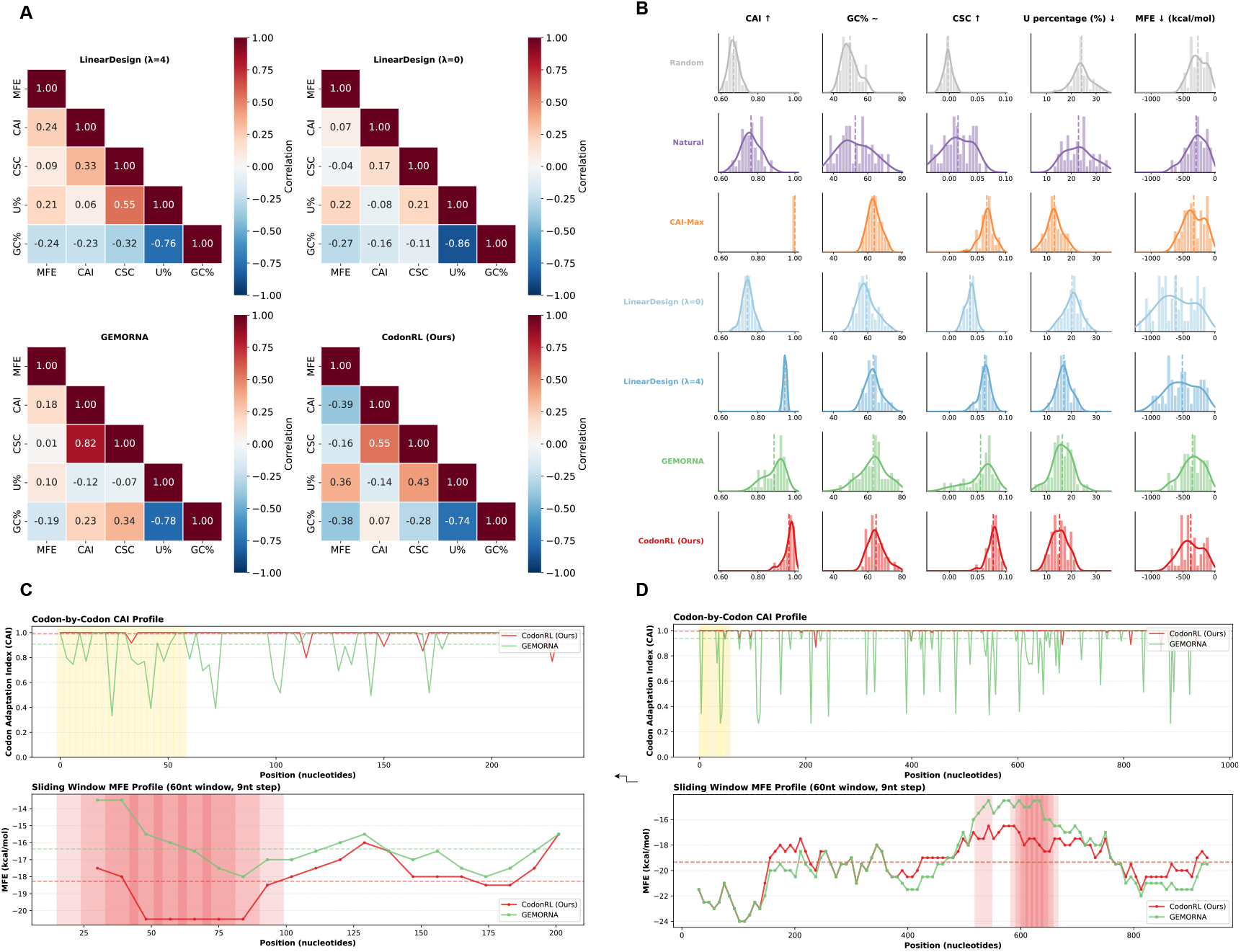
Multi-Scale benchmarking of CodonRL against state-of-the-art mRNA design methods. Spearman correlation matrices between key sequence-level objectives: minimum free energy (MFE, kcal/mol; more negative is more stable), codon adaptation index (CAI; higher predicts higher translation efficiency), codon-stability coefficient score (CSC; higher is more mRNA-stabilizing), uridine content (U%, lower is preferred for reduced innate immune activation), and GC content (GC%). Correlations are shown separately for LinearDesign (*λ* = 0), LinearDesign (*λ* = 4), GEMORNA, and CodonRL, with color indicating correlation strength (red = positive, blue = negative) and annotated values giving the correlation coefficients. Distributions of the same five metrics across all designed coding sequences for each method. Columns correspond to CAI (desired to increase), GC% (kept within a target range), CSC (desired to increase), U% (desired to decrease), and MFE (kcal/mol; more negative indicates more stable structures), and rows correspond to Random, Natural, CAI-maximization, LinearDesign (*λ* = 0), LinearDesign (*λ* = 4), GEMORNA, and CodonRL. For each method–metric pair, histograms are plotted on unified x-axis scales, with vertical dashed lines indicating the mean values (and, where applicable, top-design landmarks), highlighting that CodonRL shifts the distributions toward simultaneously higher CAI and CSC, lower U% and controlled GC%, and more favorable (more negative) MFE. **(C–D)** Codon-by-codon and local-structure profiles for two representative proteins (Glycophorin-E, shorter sequence in (C); AIMP2, longer sequence in (D)), comparing CodonRL (red) and GEMORNA (green) for the same amino-acid sequence. Top panels show codon-level CAI along the coding sequence, computed using human codon usage; shaded gold regions mark positions where CodonRL maintains or improves CAI relative to GEMORNA. Bottom panels show sliding-window MFE traces (60-nt window, 9-nt step); shaded red segments indicate windows where CodonRL achieves at least 2 kcal/mol more favorable (more negative) MFE. These illustrate that CodonRL can create mRNAs with globally improved secondary-structure stability while preserving or enhancing codon usage and transcript-stability signals relative to GEMORNA.

#### Correlation Patterns Reveal Method-Specific Trade-offs

A critical challenge in codon optimization is understanding whether different design objectives inherently conflict or can be simultaneously satisfied. To quantify this, we analyze pairwise correlations between key metrics (MFE, CAI, CSC, U%) across different methods. This analysis reveals whether a method can achieve Pareto improvement (improving multiple objectives simultaneously) or merely shifts which objectives are prioritized.

We characterized how objectives co-vary under each design strategy by computing pairwise Spearman correlations among five core metrics (MFE, CAI, CSC, U%, and GC%) for LinearDesign (*λ* = 4), LinearDesign (*λ* = 0), GEMORNA, and CodonRL (Figure 3A). We interpret these correlations relative to the desired optimization directions: higher CAI and CSC are favorable, whereas lower MFE and U% are favorable.

LinearDesign variants show stability–efficiency tension: MFE–CAI correlations are positive, meaning that stabilizing the structure (lowering MFE) is mildly associated with reduced CAI. GEMORNA moderates but does not remove this pattern. CodonRL reverses this relation, so sequences with more negative MFE now tend to have higher CAI, converting a traditional trade-off into a synergy. A similar reorientation is observed for MFE–CSC, indicating that structural improvements and codon stability become more aligned. For objectives where both directions are beneficial when they decrease, positive correlations directly represent synergy; CodonRL shows the strongest such coupling, suggesting that more stable structures are also associated with lower uridine content. At the same time, CodonRL maintains an anti-correlation between CAI and U%, so higher CAI sequences tend to have reduced uridine burden, while accepting a more pronounced trade-off between CSC and U%. Overall, CodonRL aligns structural stability with translational efficiency and uridine reduction, while concentrating remaining trade-offs into a smaller subset of metrics.

#### Metric Distributions Reveal Method-Wide Shifts in Design Behavior

Across six design strategies (Random, CAI-max, LinearDesign (*λ* = 0), LinearDesign (*λ* = 4), GEMORNA, and CodonRL), the distributions of CAI, MFE, CSC, GC%, and U% reveal distinct optimization priorities and trade-offs (Figure 3B). Random sampling produces broad, unstructured distributions across metrics. CAI-max concentrates mass at high CAI values but degrades structural and compositional balance. LinearDesign (*λ* = 0) favors low MFE values but yields only moderate CAI and CSC, whereas *λ* = 4 partially restores this balance. GEMORNA shifts the distributions toward a more even profile, increasing CAI and CSC while maintaining reasonable structural stability.

CodonRL achieves the most favorable global configuration: its CAI and CSC distributions peak at higher values, its MFE distribution is shifted toward more stable sequences, its GC% profile is narrow and well-centered, and its U% distribution is skewed toward lower values. Together, these distributions show that CodonRL occupies desirable regions of the design space that other methods only partially access.

#### Local Structure and Codon-Level Effects Support Global Improvements

To assess codon-level behavior, we report per-codon CAI profiles and sliding-window MFE (60-nt window, 9-nt step) for two representative proteins of different lengths, highlighting how local translation efficiency and folding stability vary across methods (Figure 3C–D).

Across both proteins, CodonRL maintains uniformly higher CAI profiles relative to GEMORNA across substantial portions of the sequence, highlighted by extended regions where CodonRL’s per-codon CAI is consistently superior. The MFE traces show similarly broad segments where CodonRL achieves more favorable local folding stability.

### Multi-objective tradeoffs

CodonRL enables user-directed tradeoffs between multiple objectives using weights *α*_*o*_ for each objective *o* at inference time. To examine how weighting during inference controls multi-objective codon optimization, and to test the flexibility of our two-stage framework, we performed systematic sweeps over the scalar weights *α*_CAI_ and *α*_U_, which modulate the relative importance of translation efficiency and uridine minimization, respectively (Figures 4 and 5).

**Figure 4:**
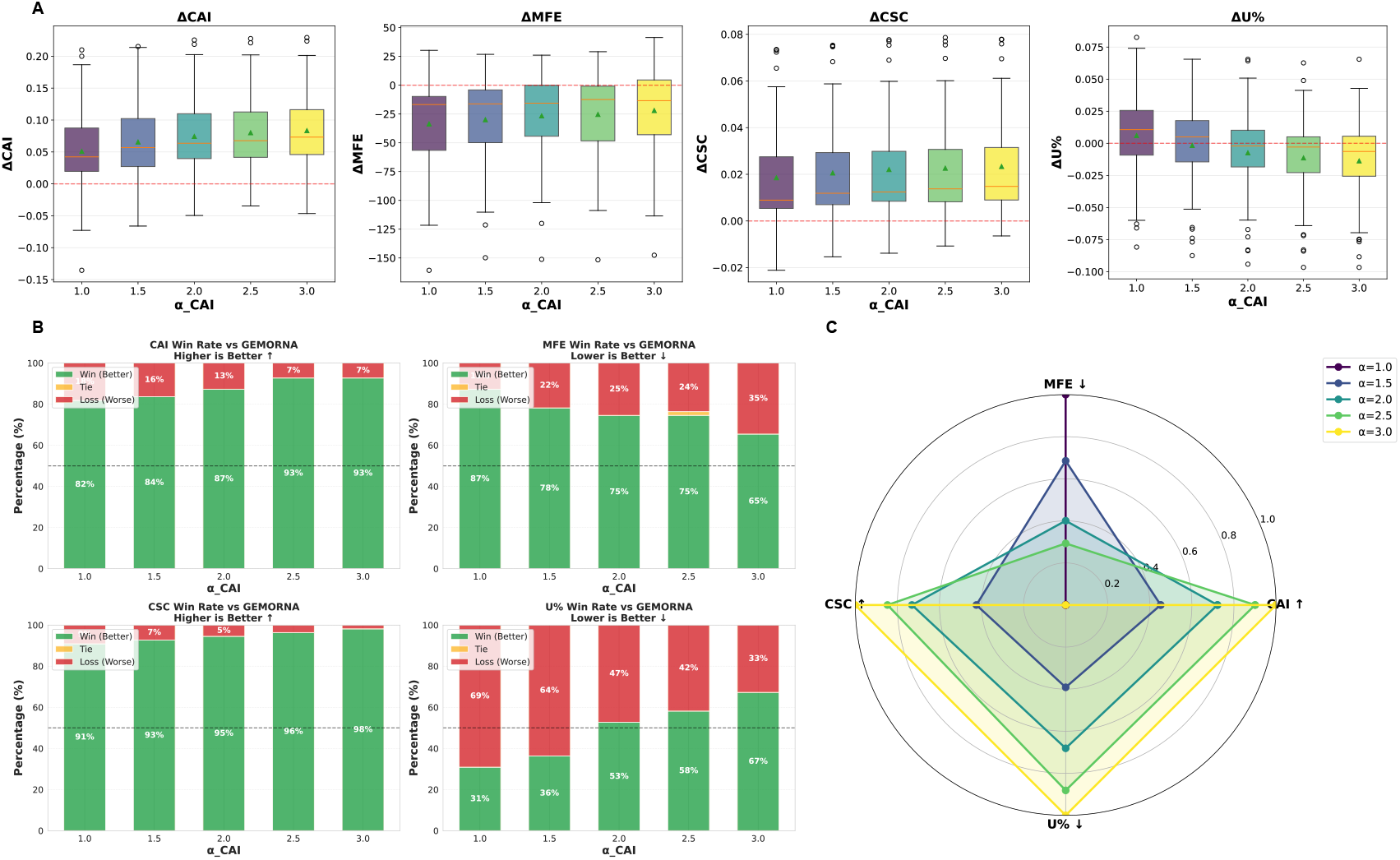
Multi-objective evaluation of *α*-weighted codon optimization across CAI, MFE, CSC, and U-content. **(A)** Box-and-whisker plots show per-sequence changes in codon adaptation index (ΔCAI), minimum free energy (ΔMFE), codon stabilization coefficient (ΔCSC), and uridine fraction (Δ*U* %) relative to the GEMORNA baseline across *α*_CAI_ ∈ { .0, 1.5, 2.0, 2.5, 3.0} . Individual points denote sequences and colored triangles denote group means. Higher ΔCAI and ΔCSC indicate improved codon usage and predicted mRNA stability, whereas lower ΔMFE and Δ*U* % reflect enhanced structural stability and reduced uridine burden. Increasing *α*_CAI_ yields monotonic gains in ΔCAI and ΔCSC, with moderate trade-offs in MFE and a mild increase in U-content at high *α*. **(B)** Stacked bar charts report win–tie–loss rates against GEMORNA for each *α*_CAI_ across four metrics: CAI (higher is better), MFE (lower is better), CSC (higher is better), and U% (lower is better). Bars quantify, for each metric, the fraction of proteins for which the optimized sequences outperform (green), match (yellow), or underperform (red) the baseline. **(C)** Radar plot summarizing normalized performance trends across the four metrics as a function of *α*_CAI_. Higher radial extent indicates stronger improvement for metrics where higher values are better (CAI, CSC), and stronger reduction for metrics where lower values are favorable (MFE, U%). The plot highlights coordinated gains in CAI and CSC as *α*_CAI_ increases, accompanied by predictable structural and compositional trade-offs.

**Figure 5:**
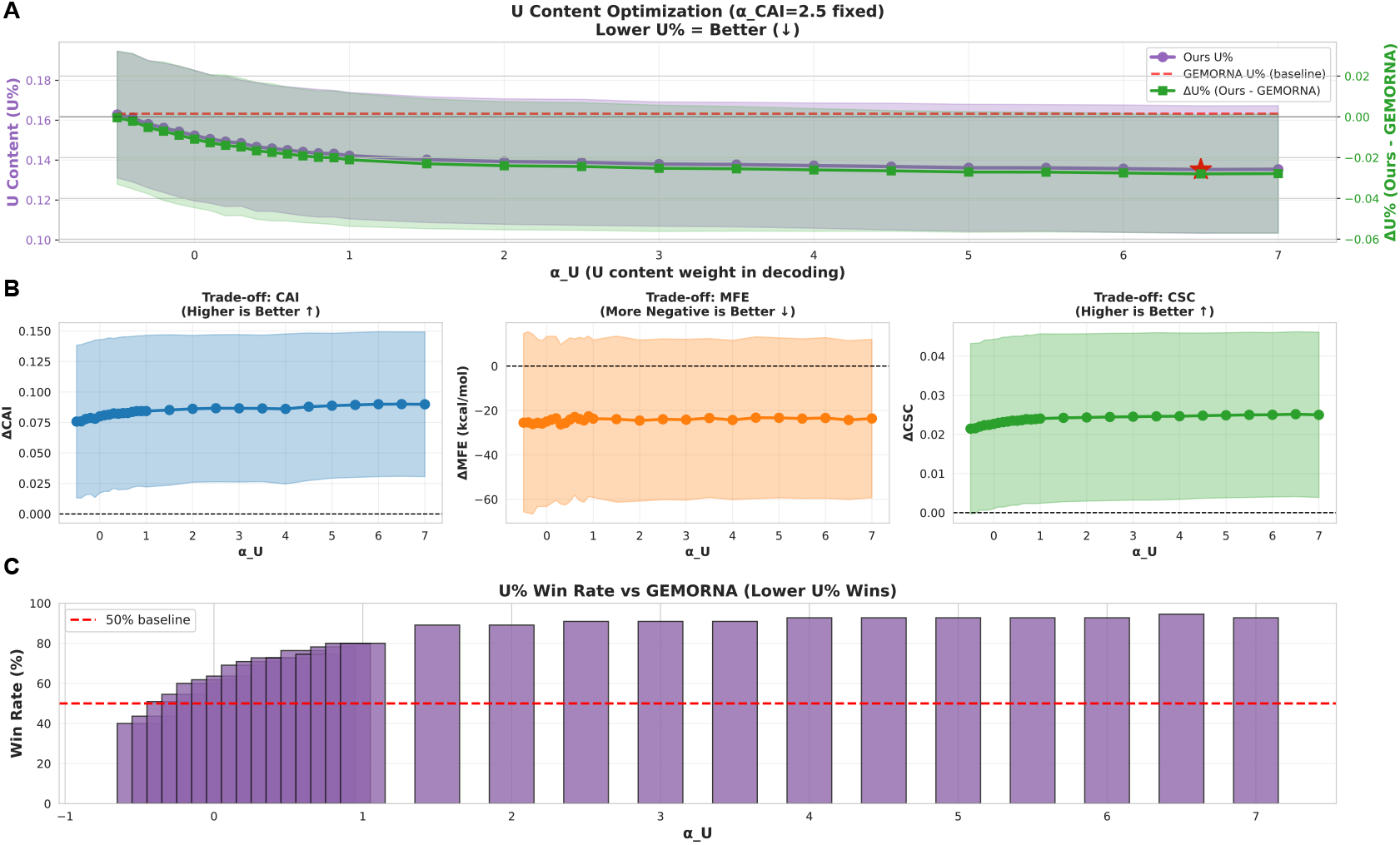
U content optimization through *αU* weight tuning with fixed *αCAI* = 2.5. **(A)** U content (U%) optimization landscape showing the effect of *αU* on uridine percentage in designed mRNA sequences. Purple line with shaded region indicates ± mean standard deviation of U% in our method across different *αU* values. Red dashed line represents the baseline U% from GEMORNA. Green line (right y-axis) shows ΔU% (Ours - GEMORNA), with the red star marking the optimal *αU* = 6.5 that achieves minimal U content while maintaining structural stability. Lower U% is desirable for reduced immunogenicity. **(B)** Trade-off analysis between U content reduction and other mRNA design objectives. Left panel: ΔCAI (Codon Adaptation Index) remains positive across all *αU* values, indicating preserved translation efficiency. Middle panel: ΔMFE (Minimum Free Energy) shows consistent negative values around -25 kcal/mol, demonstrating maintained thermodynamic stability. Right panel: ΔCSC (Codon Stabilization Coefficient) maintains positive values, confirming preservation of codon pair optimization. **(C)** Win rate comparison against GEMORNA baseline across the *αU* sweep. Bar heights represent the percentage of sequences where our method achieves lower U% than GEMORNA. Win rates consistently exceed 50% baseline for *αU ≥* 0, with peak performance above 90% win rate for *αU≥* [2, 7], demonstrating robust superiority in U content minimization while balancing multiple design constraints.

**Figure 6:**
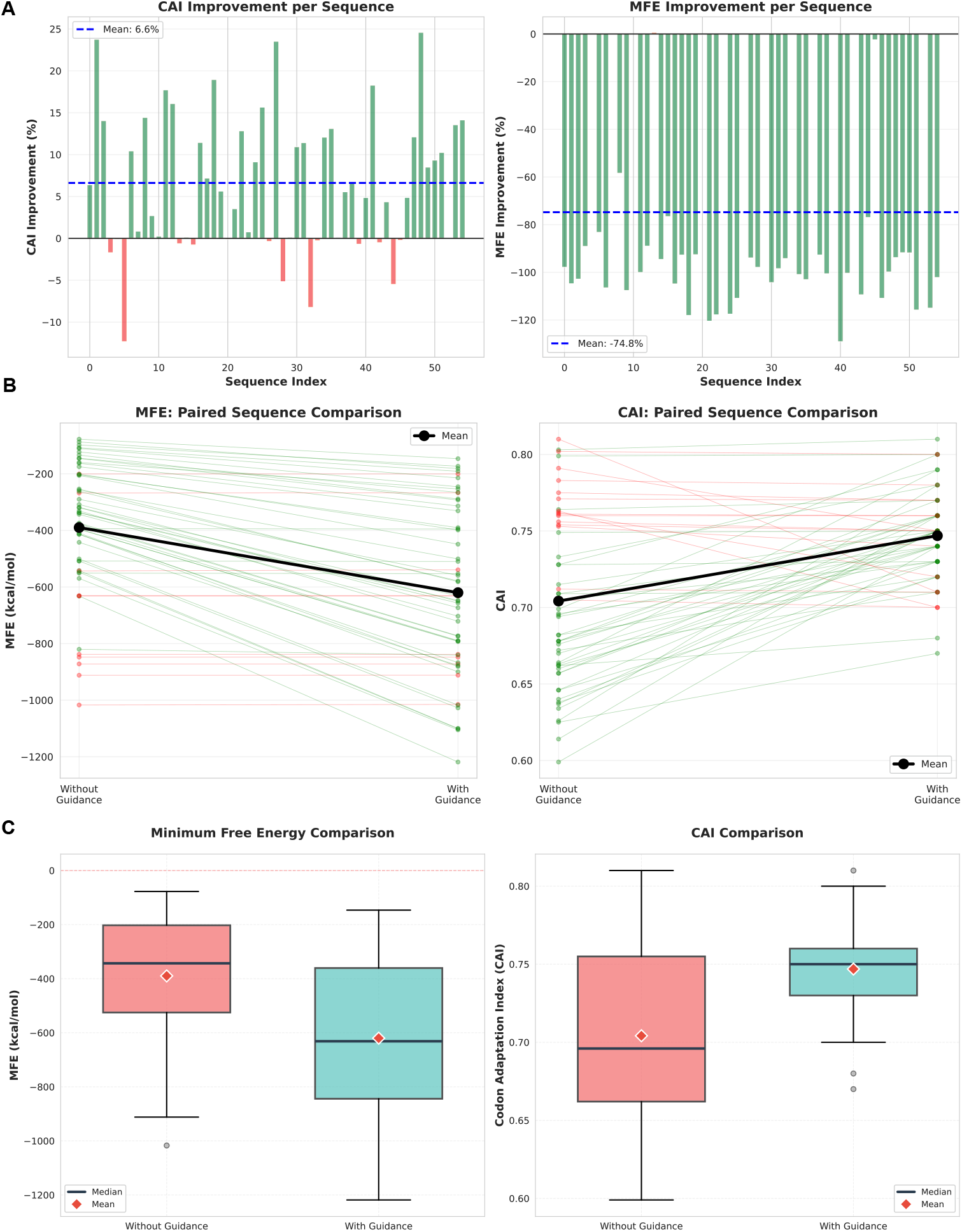
Impact of demonstration-guided learning on codon optimization performance during training. **(A)** Per-sequence improvement percentages for CAI (left) and MFE (right) with guidance relative to without guidance. Green bars indicate improvements while red bars indicate degradation. Dashed lines show mean improvements for CAI and MFE (more negative MFE indicates more stable structure). **(B)** Paired comparison of individual sequences showing trajectory changes from without guidance (left) to with guidance (right) for MFE (left panel) and CAI (right panel). Each line represents one sequence, with green lines indicating improvement and red lines indicating degradation after applying guidance. Black bold lines show mean values. **(C)** Distribution comparison via box plots for MFE (left) and CAI (right) between without guidance (coral) and with guidance (teal). Guidance significantly improves both MFE stability and CAI. dynamic adjustment of design priorities and facilitates adaptation to diverse optimization criteria. Empirical evaluation on a diverse set of proteins demonstrates consistent improvements across MFE, CAI, CSC, and uridine-related metrics, with performance surpassing the previous state-of-art method. Overall, the framework establishes a scalable and adaptable foundation for codon optimization under complex and variable biological constraints.

#### Monotonic CAI Gains with Predictable Trade-offs

Increasing *α*_CAI_ ∈ {1.0, 1.5, 2.0, 2.5, 3.0} (with all other weights fixed) yields monotonic gains in translation efficiency relative to GEMORNA, with ΔCAI increasing steadily and ΔCSC rising in parallel (Figure 4A).

The associated structural and compositional trade-offs remain modest. As *α*_CAI_ increases, ΔMFE becomes slightly less negative, indicating a small relaxation of stability gains, while U% shows minimal drift except at the highest *α*_CAI_ value (Figure 4A). Together, these trends indicate that CAI improvements can be emphasized without substantially disrupting the overall design profile.

Consistent with these shifts, CAI win rates against GEMORNA increase monotonically with *α*_CAI_, while MFE remains strong across all settings and CSC maintains high win rates with few losses (Figure 4B). U% performance also improves with *α*_CAI_, with win rates rising from below 50% at lower settings to above 50% at higher settings.

Overall, varying *α*_CAI_ produces a smooth and controllable trade-off surface: improvements in CAI and CSC strengthen with increasing weight, whereas gains in MFE and U% contract only modestly (Figure 4C).

#### Uridine Minimization and High-Confidence Control with Minimal Interference

Sweeping *α*_U_ *∈* [*»*1, 7] with *α*_CAI_ fixed at 2.5 yields monotonic reductions in uridine fraction (U%), reaching the lowest values near *α*_U_ = 6.5 (Figure 5A). Relative to GEMORNA, ΔU% remains negative across most of the sweep and plateaus for *α*_U_ *∈* [5, 7], indicating robust improvements in uridine minimization. Notably, U% is already reduced at *α*_U_ = 0, suggesting that the learned policy implicitly disfavors uridine-rich compositions.

These gains show minimal interference with other objectives (Figure 5B). Across the sweep, ΔCAI remains positive, ΔMFE remains consistently negative with only narrow fluctuations, and ΔCSC remains positive and stable, indicating preserved translation efficiency, maintained structural stability, and retained codon-stability preferences. Together, these trends suggest that U% minimization is largely orthogonal to the other objectives over the explored range.

Consistent with these improvements, the fraction of proteins for which CodonRL achieves lower U% than GEMORNA exceeds 50% for all *α*_U_ *≥* 0 and rises above 90% for *α*_U_ *∈* [2, 7] (Figure 5C), indicating a broad parameter regime that reliably produces low-uridine designs without compromising other metrics.

#### Design Space Coverage and the Learned Pareto Frontier

CodonRL exhibits a smooth and predictable response to weight tuning. The frozen CodonRL supplies a strong structural prior, enabling shifts in objective emphasis without retraining. The resulting performance envelopes (Figures 4 and 5) demonstrate that a single model can traverse the CAI–MFE–CSC–U% design space through continuous *α*-adjustments. This train-once, tune-at-inference approach contrasts with pipelines such as LinearDesign or GEMORNA, whose objective coupling limits flexible multi-objective navigation.

### Impact of Demonstration-Guided Learning

A key innovation of CodonRL is the integration of expert demonstrations to accelerate policy learning and improve sample efficiency. To quantify the contribution of this design choice, we conducted an ablation study comparing CodonRL trained with and without demonstration-guided pre-population of the replay buffer (Figure 6).

#### Demonstration Guidance Substantially Improves MFE Optimization

Including expert demonstrations improves performance relative to cold-start training across most proteins, with the largest gains observed for MFE (Figure 6A). In particular, MFE becomes more favorable (more negative) for the majority of sequences, indicating that demonstration guidance effectively bootstraps learning for global structure optimization, where unguided exploration rarely discovers stable folding configurations.

Improvements in CAI are smaller but remain consistently positive across most sequences (Figure 6A). This asymmetry reflects the different optimization difficulty of the objectives: CAI is comparatively local and often amenable to greedy improvements, whereas MFE depends on coordinated codon choices across the full sequence to create favorable long-range base-pairing patterns. Together, these results show that demonstration-guided learning is particularly valuable for the more challenging structural optimization objective.

#### Paired Trajectory Analysis Reveals Consistent Learning Enhancement

Paired per-protein comparisons show that demonstration guidance improves both structural stability and translation efficiency across the benchmark set (Figure 6B). With guidance, MFE becomes more negative for the majority of sequences, indicating more stable predicted structures, and CAI increases for most sequences, indicating improved translation efficiency. The mean effect is consistent with these per-protein shifts, supporting a broad benefit of guidance rather than improvements driven by a small subset of proteins.

These paired comparisons reveal that demonstration guidance does not simply improve average performance through a few outlier sequences, but rather provides consistent benefits across the entire benchmark set. The guidance mechanism successfully transfers structural optimization knowledge from LinearDesign’s dynamic programming solutions to the RL policy, enabling the learned model to combine expert-level structural design with the flexibility of reinforcement learning’s multi-objective framework.

#### Distribution Analysis Confirms Systematic Benefits

Distribution-level comparisons confirm that demonstration guidance improves stability and increases consistency across proteins (Figure 6C). With guidance, MFE shifts toward more negative values, with improved medians and interquartile ranges, indicating that stability gains apply to most sequences rather than a small subset. CAI also increases in median and mean, although the distributions overlap more strongly, consistent with CAI being less dependent on long-range coordination than MFE. Finally, guidance reduces variability across proteins, suggesting that replay-buffer pre-population stabilizes training outcomes.

## Conclusion

We introduced a reinforcement learning framework for multi-objective codon sequence optimization that integrates expert demonstrations, efficient structural feedback, and an efficient transformer-based Q-network. By combining LinearFold for rapid intermediate evaluation with ViennaRNA for accurate final assessment, the method reduces folding-related computational cost while maintaining biological fidelity. Demonstration pre-population further accelerates training and enables effective use of optimized sequences within the learning process. More importantly, the trained model supports flexible inference, allowing different metric definitions or objective functions to be incorporated at generation time without retraining. This capability enables dynamic adjustment of design priorities and facilitates adaptation to diverse optimization criteria. Empirical evaluation on a diverse set of proteins demonstrates consistent improvements across MFE, CAI, CSC, and uridine-related metrics, with performance surpassing the previous state-of-art method. Overall, the framework establishes a scalable and adaptable foundation for codon optimization under complex and variable biological constraints.

## Methods

CodonRL is a reinforcement learning-based framework for mRNA sequence design that addresses the combinatorial challenge of codon selection while optimizing multiple competing objectives. Figure 1 provides a comprehensive overview of CodonRL. An introduction to reinforcement learning and the complete algorithmic specifications of CodonRL are provided in the Supplementary Material (Section A).

### The Codon Optimization Challenge

The genetic code’s degeneracy creates an exponentially large search space for mRNA design (Figure 1C). For a protein of length *L* amino acids, where each amino acid cank be encoded by *k*_*i*_ *∈* [1, 6] synonymous codons, the total number of possible mRNA sequences is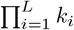. With average codon degeneracy *k ∈* [3, 6], this yields approximately *k*^*n*^ possible sequences. For a typical therapeutic protein of 300 amino acids, this translates to *∼* 10^180^ possible mRNA sequences—far exceeding the number of atoms in the observable universe. This combinatorial explosion necessitates intelligent optimization strategies rather than exhaustive enumeration.

### Two-Stage Learning Framework

Our approach decouples policy learning from objective optimization through a two-stage framework (Figure 1A–B). In the **training stage** (Figure 1A), we employ reinforcement learning to train a transformer-based policy network that learns to generate structurally stable mRNA sequences through interactions with a thermodynamic reward signal. The architecture consists of a protein encoder that processes the target amino acid sequence into contextualized representations, and an autoregressive codon decoder that outputs Q-values for all possible codons at each position *t*. The policy is trained via *ϵ*-greedy exploration with experience replay, optimizing the Bellman equation through temporal difference learning.

In the **inference stage** (Figure 1B), we freeze the trained policy network and augment its Q-values with weighted objective functions to achieve multi-objective optimization. This frozen policy serves as a learned prior that encodes structural stability preferences, while tunable hyperparameters *{α*_CAI_, *α*_CSC_, *α*_GC_, *α*_U_*}* allow flexible control over translation efficiency (CAI), codon pair stability (CSC), GC content, and uridine content. We define the state at position *t* as *s*_*t*_ = (**H**_protein_, *c*_1:*t−*1_), where **H**_protein_ is the cached protein encoder output for the target amino acid sequence of length *L*, and *c*_1:*t−*1_ = (*c*_1_, *c*_2_, …, *c*_*t−*1_) is the sequence of codons selected at all previous positions. The final codon selection combines learned Q-values with objective scores:

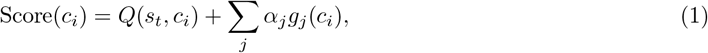

where *Q*(*s*_*t*_, *c*_*i*_) represents the learned Q-value for codon *c*_*i*_ given state *s*_*t*_ (protein context and history), and *g*_*j*_(*c*_*i*_) are deterministic objective functions with weights *α*_*j*_.

### Architectural Components

#### Multi-Objective Reward Design

We introduce a multi-objective reward formulation for the training stage that addresses trade-offs in codon optimization:

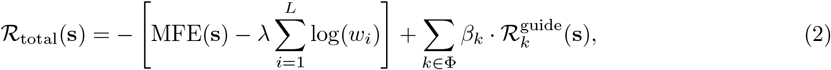

where **s** represents the mRNA sequence, *w*_*i*_ denotes the relative adaptiveness of the *i*-th codon in the sequence defined as *w*_*i*_ = *f*_*i*_*/* max_*c∈*syn(*i*)_ *f*_*c*_ with *f*_*i*_ being the codon frequency in the target organism’s usage table and syn(*i*) representing all synonymous codons for the same amino acid, *β*_*k*_ represents the milestone reward weight (default 0.3) applied at position *k*, and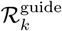 represents the milestone-specific reward calculated as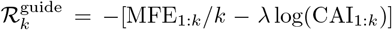 for partial sequence up to position *k*, and Φ = {*⌊*0.25*L⌋, ⌊*0.5*L⌋, ⌊*0.75*L⌋*} denotes the milestone positions as fractions of the total protein length *L*.

This reward function balances three critical objectives: (i) the relative adaptiveness of codons based on species-specific translation efficiency metrics, (ii) an empirically optimized trade-off parameter *λ* that mediates between mRNA stability (MFE) and codon adaptation index (CAI), and (iii) milestone rewards at quartile completion points that densify the reward signal and accelerate convergence.

#### Efficient Structural Feedback

To address the computational bottleneck of RNA secondary structure prediction while maintaining biological accuracy, we propose a hierarchical folding evaluation strategy that leverages two complementary algorithms. LinearFold provides *O*(*L*) complexity predictions at intermediate milestones, enabling rapid feedback during training, while ViennaRNA performs thermodynamically rigorous calculations for final sequence evaluation. This approach reduces overall computational cost without sacrificing prediction accuracy (Pearson correlation above 0.9 between LinearFold and ViennaRNA [Huang et al., 2019]). Our benchmarking experiments demonstrate that LinearFold achieves substantial speedup over ViennaRNA while maintaining prediction quality (Supplementary Figure 7).

#### Cached Transformer Architecture for Q-Network

We present a neural architecture that combines the representational power of efficient flash attention with the decision-making capabilities of Q-learning, specifically designed for coding sequence optimization.

##### Protein Encoder Caching

The protein encoder computes contextualized representations once per episode and caches them for reuse:

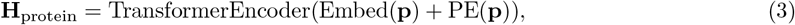

where **p** represents the input protein sequence, Embed(*·*) denotes the token embedding layer, and PE(*·*) is the positional encoding. This caching mechanism eliminates redundant computations across the *L* sequential decisions, reducing per-episode complexity from *O*(*L*^3^) to *O*(*L*^2^).

##### Flash Attention-based Deep Q-Learning

We integrate Flash Attention 2 (see Supplementary Section A for detailed review) into our transformer-based Q-network, achieving linear memory complexity *O*(*L*) instead of quadratic *O*(*L*^2^) for sequence length *L*. The complete architecture consists of a transformer encoder for the protein sequence with model dimension *d*_model_ = 512 and 8 attention heads, coupled with a transformer decoder for the mRNA sequence. The decoder employs cross-attention mechanisms to attend to the cached protein representations, enabling the model to dynamically focus on relevant amino acid context during codon selection. All layers use position-wise feed-forward networks with inner dimension 2048, following standard transformer design principles while maintaining computational efficiency through Flash Attention’s optimized attention kernel.

##### Position-aware Q-value Generation

Our architecture generates position-specific Q-values by attending to both local decoder context at position *t* and the global protein structure:

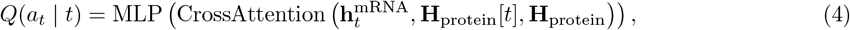

where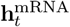 is the decoder hidden state at position *t*, **H**_protein_[*t*] denotes the protein encoder representation of the corresponding amino acid at position *t*, and **H**_protein_ is the full protein encoder output (used as keys and values). This formulation enables action-value estimation conditioned on both the local amino acid identity and the global protein sequence context.

#### Training Procedure

To accelerate early-stage learning and improve sample efficiency, we introduce a replay buffer warm-start strategy using high-quality guide trajectories from expert-designed sequences, such as those produced by LinearDesign:

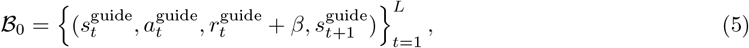

where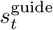 and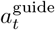 are the state and action at time *t* in the guide sequence, 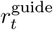 is the corresponding reward, and *β* = 0.1 provides an auxiliary bonus for imitation of demonstrated trajectories. This pre-population bootstraps the Q-function with informative examples before online exploration begins.

We further enhance training efficiency through asynchronous MFE calculation, hierarchical caching, and mixed precision training (see Supplementary Section A for details).

#### Inference Procedure

At inference time, we employ a flexible multi-objective decoding strategy that extends beyond the training objectives to accommodate diverse optimization criteria. Given a trained Q-network and a target protein sequence, our inference procedure generates optimized mRNA sequences through a position-wise greedy selection process with composite scoring.

##### Multi-Objective Decoding Framework

For each position *t* in the protein sequence, we compute a composite score for each candidate codon *c* that combines the learned Q-values with additional objective-specific terms:

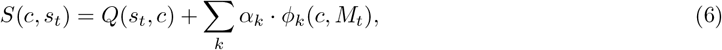

where *Q*(*s*_*t*_, *c*) represents the Q-value from the trained network, *α*_*k*_ denotes objective-specific weights, and *ϕ*_*k*_(*c, M*_*t*_) represents feature functions computed over the candidate codon *c* and partially constructed sequence *M*_*t*_. Eq. 6 restates Eq. 1 with explicit state dependence: Score(*c*) *≡ S*(*c, s*_*t*_) and *g*_*k*_(*c*) is a special case of *ϕ*_*k*_(*c, M*_*t*_) when the feature is prefix-independent.

##### Objective-Specific Features

We tested the following optimization objectives during inference:

- **Codon Adaptation Index:** *ϕ*_CAI_(*c*) = log(*w*_*c*_) where *w*_*c*_ is the relative adaptiveness of codon *c*.
- **mRNA Stability:** *ϕ*_CSC_(*c*) = log(CSC_*c*_) where CSC_*c*_ is the codon stabilization coefficient.
- **GC Content:** *ϕ*_GC_(*c, M*_*t*_) = *−*|*g*(*M*_*t*_ *∪ {c}*) *− g*_target_| where *g*(*·*) computes GC content and *g*_target_ is the desired GC level.
- **Uridine Content:** *ϕ*_U_(*c*) = *−u*(*c*) to directly minimize uridine usage, where *u*(*·*) computes the proportion of uridine nucleotides in the input sequence.

The GC and uridine content features enable dynamic trajectory adjustment by evaluating how each candidate codon affects the cumulative sequence composition.

##### Decoupled Training and Inference Objectives

An advantage of our approach is the decoupling of training and inference objectives. While the Q-network is trained to optimize CAI and MFE, the inference procedure can flexibly incorporate additional or alternative objectives (such as CSC, GC content, or uridine minimization) without retraining. This is achieved by treating the learned Q-values as a foundation that captures general codon optimization principles, which are then augmented with objective-specific terms during decoding.

##### Computational Efficiency

The inference procedure maintains the computational advantages of our architecture through protein encoder caching. The protein representation **H**_protein_ is computed once per sequence and reused across all *L* decoding steps, requiring only *O*(*L*^2^) total computation rather than *O*(*L*^3^). At each decoding position *t*, we evaluate a set of candidate codons corresponding to the target amino acid. Let *A*_*t*_ denote the set of synonymous codons for the amino acid at position *t*, with cardinality |*A*_*t*_| (typically 2–6). This makes the per-position scoring complexity *O*(|*A*_*t*_|) after the cached attention mechanism.

## Acknowledgments

This work was supported in part by the US National Science Foundation [III-2232121], the US National Institutes of Health [R01HG012470]. S.D. was partially supported by a SoftBank Group–Arm Fellowship. Conflict of Interest: C.K. is a co-founder of Ellumigen, Inc.

## A Supplementary Material

### Reinforcement Learning Background

Reinforcement learning provides a principled mathematical framework for sequential decision-making under uncertainty. In this paradigm, an agent interacts with an environment by selecting actions based on observed states, receiving scalar reward signals that quantify the immediate quality of each decision [Li, 2017]. The fundamental objective is to learn a policy that maximizes the expected cumulative discounted reward over time, balancing short-term gains against long-term consequences. This framework naturally accommodates codon optimization, where each codon selection influences not only immediate sequence properties but also downstream optimization potential through effects on secondary structure and compositional constraints.

Q-learning constitutes a canonical value-based reinforcement learning algorithm that learns an action-value function representing the expected long-term reward of executing a specific action in a given state and subsequently following the optimal policy. The Q-function satisfies the Bellman optimality equation, which forms the basis for iterative learning through temporal difference updates. In deep Q-learning, the Q-function is approximated by a neural network with parameters *θ*, enabling application to high-dimensional state and action spaces. The network is trained by minimizing the temporal difference error between predicted Q-values and target values computed using the Bellman equation, gradually refining its estimates as more experience is accumulated.

We formulate codon optimization as a sequential decision problem amenable to reinforcement learning. In this framework, an agent learns to construct optimal mRNA sequences by iteratively selecting codons for each amino acid position, with the objective of maximizing long-term sequence quality. At each position, the agent observes a state representation encoding the partial mRNA sequence and protein context, then selects an action corresponding to a synonymous codon choice. The quality of this decision is quantified by a reward signal that evaluates structural stability and translation efficiency. A neural Q-network learns to estimate the expected cumulative reward for each possible codon selection, guiding the agent toward sequences that balance multiple competing objectives. The learning process leverages experience replay, where past decision trajectories are stored in a buffer and resampled during training to improve sample efficiency and stability. Unlike supervised learning approaches that require extensive experimental labels, reinforcement learning enables the discovery of effective optimization strategies through iterative exploration guided by computationally tractable objective functions.

### Additional Related Work

#### Codon Optimization Methods

Codon optimization has evolved from simple frequency-matching approaches to sophisticated algorithms incorporating multiple biological constraints. Early methods focused exclusively on CAI maximization [Sharp and Li, 1987], replacing rare codons with frequently used synonymous alternatives. However, these approaches often resulted in decreased expression due to mRNA instability or translational bottlenecks.

The recognition of mRNA secondary structure’s importance led to structure-aware optimization methods. LinearDesign [Zhang et al., 2023] represented a significant advance, formulating codon optimization as a joint problem of maximizing CAI while minimizing MFE through dynamic programming, achieving up to 128-fold improvement in COVID-19 vaccine antibody responses. While effective, LinearDesign employs a beam search algorithm that makes locally optimal decisions without the ability to explore radically different solution paths.

Recent work has explored machine learning approaches to codon optimization. CodonTransformer [Fallahpour et al., 2025] employs transformer architectures with STREAM encoding trained on over 1 million DNA-protein pairs, while ICOR [Jain et al., 2023] uses bidirectional LSTM networks to capture sequential context. These supervised approaches require extensive experimental data.

#### Reinforcement Learning in Biological Sequence Design

Reinforcement learning (RL) has demonstrated remarkable success in biological sequence design problems. Yang et al. [2025] applied RL to regulatory element design, achieving superior performance in generating high-fitness promoters and enhancers. For protein engineering, Wang et al. [2023] developed EvoPlay using self-play RL with Monte Carlo tree search, achieving 7.8-fold bioluminescence improvement in luciferase engineering, while Sun et al. [2025] combined deep learning mutational prediction with RL for *β*-lactamase optimization.

However, codon optimization presents unique challenges not addressed in previous work: maintaining amino acid sequence identity, multi-objective optimization, and alleviating the computational expense of repeated RNA folding evaluations.

#### Efficient Transformer Architectures

Transformers have revolutionized sequence modeling but suffer from quadratic memory and computational complexity. Recent advances in efficient attention mechanisms, particularly Flash Attention [Dao et al., 2022] and Flash Attention 2 [Dao, 2023], enable linear memory scaling through tiled computation, crucial for handling long sequences in codon optimization (often exceeding 3000 nucleotides).

The application of transformers to biological sequences has shown remarkable success. ESM-1b [Rives et al., 2021] demonstrated that protein language models learn structural and functional properties from sequence alone, while the Nucleotide Transformer [Dalla-Torre et al., 2025] provides foundation models for genomic analysis. Our work builds on these foundations, adapting transformer architectures specifically for action-value estimation in Q-learning while incorporating domain-specific optimizations such as protein encoder caching.

### Computational Efficiency Optimizations

#### Asynchronous MFE Calculation

We develop an asynchronous computation pipeline that overlaps neural network forward passes with RNA folding calculations, using a thread pool executor with *n* = 4 workers. This parallelization reduces wall-clock training time compared to synchronous evaluation.

#### Hierarchical LRU Caching

We employ a two-tier Least Recently Used (LRU) caching strategy with 8,192 entries for LinearFold calculations and 4,096 entries for ViennaRNA computations.

#### Mixed Precision Training

We leverage PyTorch’s automatic mixed precision (AMP) training with the default GradScaler to accelerate computation on modern GPUs. Our implementation combines FP16 computations with FP32 master weights, achieving memory reduction while maintaining numerical stability through automatic loss scaling.

### Detailed Algorithms

Algorithm 1 gives the top-level training loop. It optionally warm-starts the replay buffer using expert trajectories (Algorithm 7), selects actions with constrained *ϵ*-greedy decoding (Algorithm 3), computes rewards using milestone densification and terminal evaluation (Algorithms 6 and 5), and updates the Q-network by Bellman optimization with experience replay (Algorithm 4). Algorithm 2 describes inference-time multi-objective decoding using the composite scoring function. The CodonRL forward pass used by both stages is summarized in Algorithm 8.

#### Algorithm 1

CodonRL Training Stage

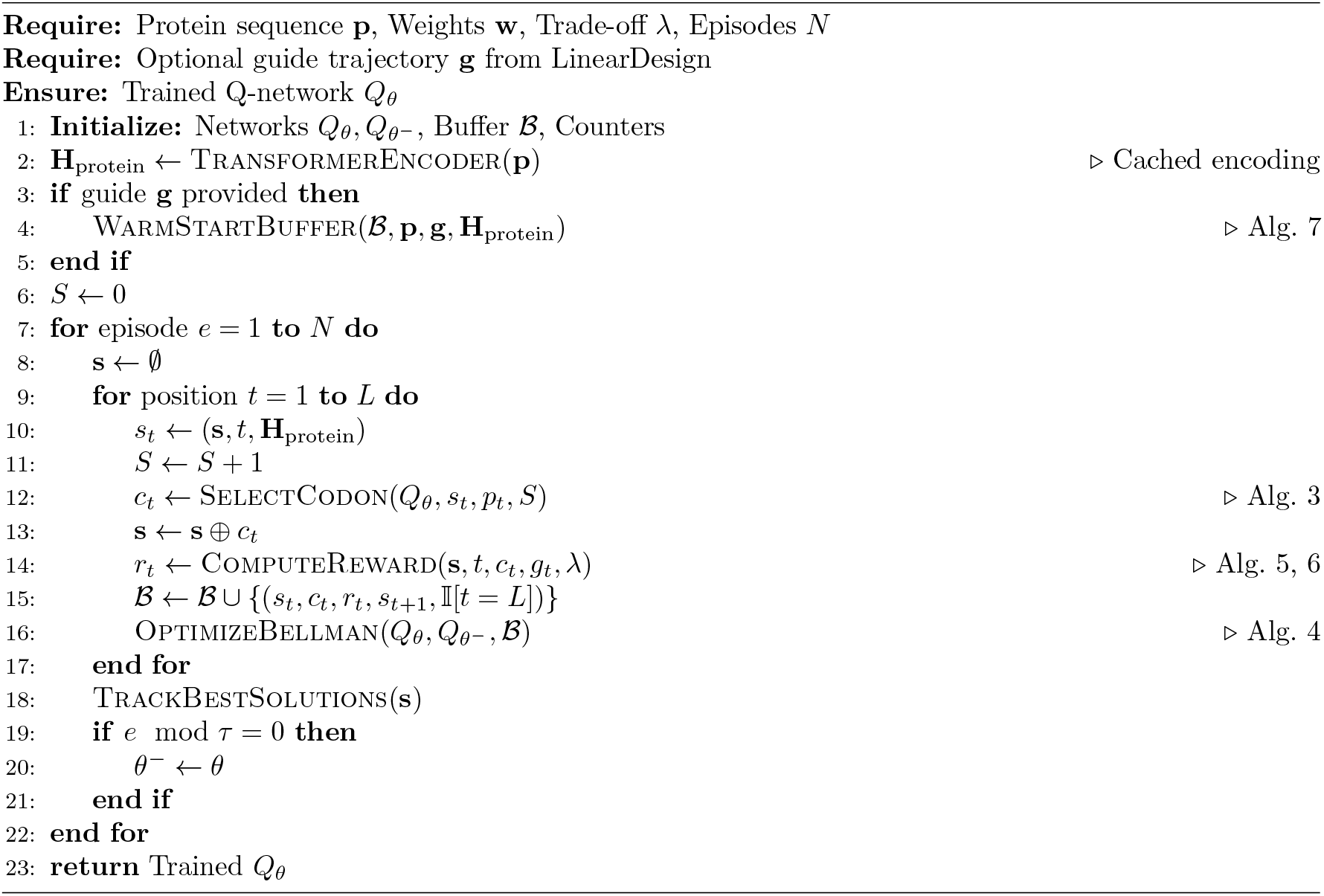

#### Algorithm 2

CodonRL Inference Stage: Multi-Objective Decoding

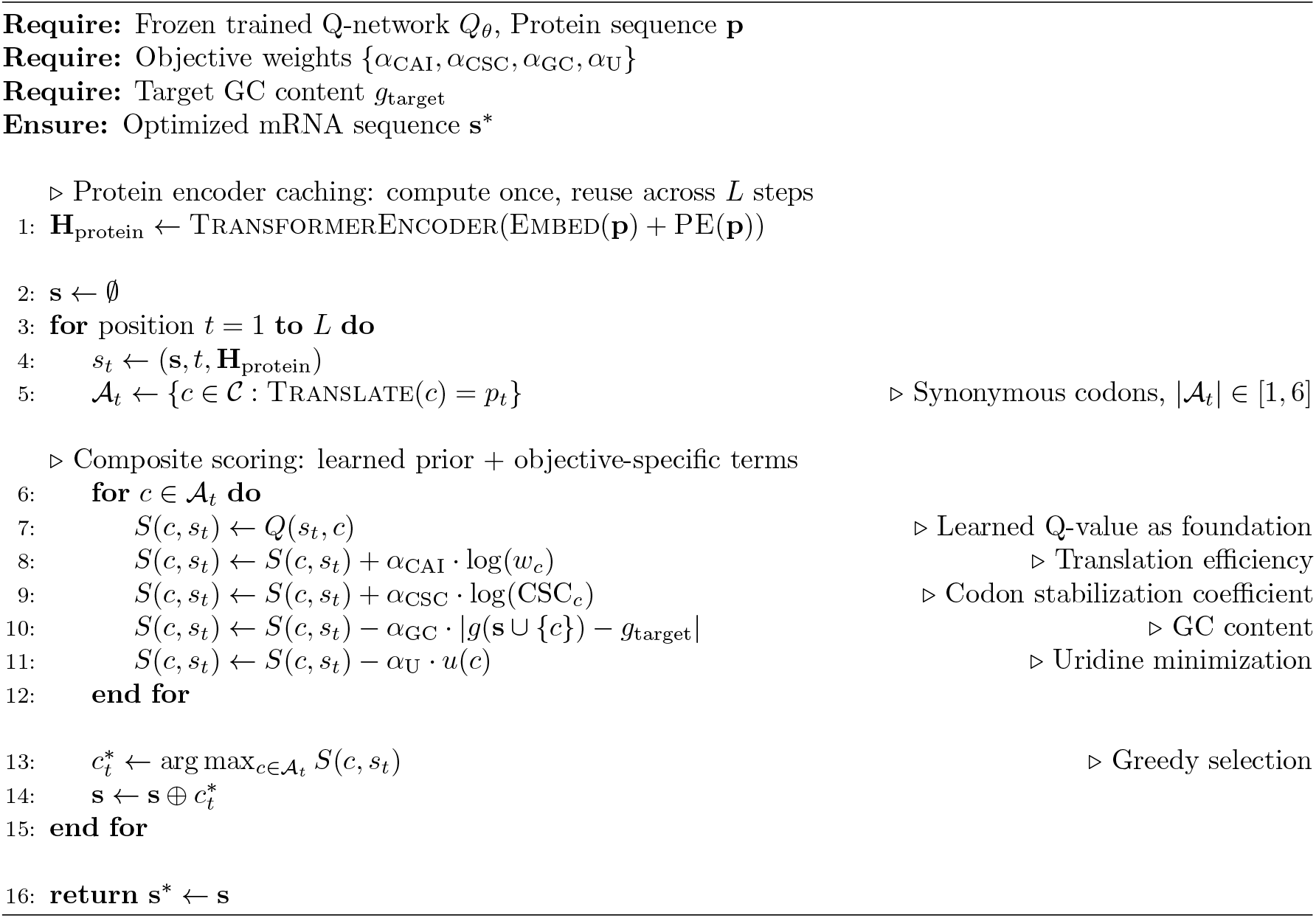

#### Algorithm 3

Constrained ϵ-Greedy Action Selection (SelectCodon)

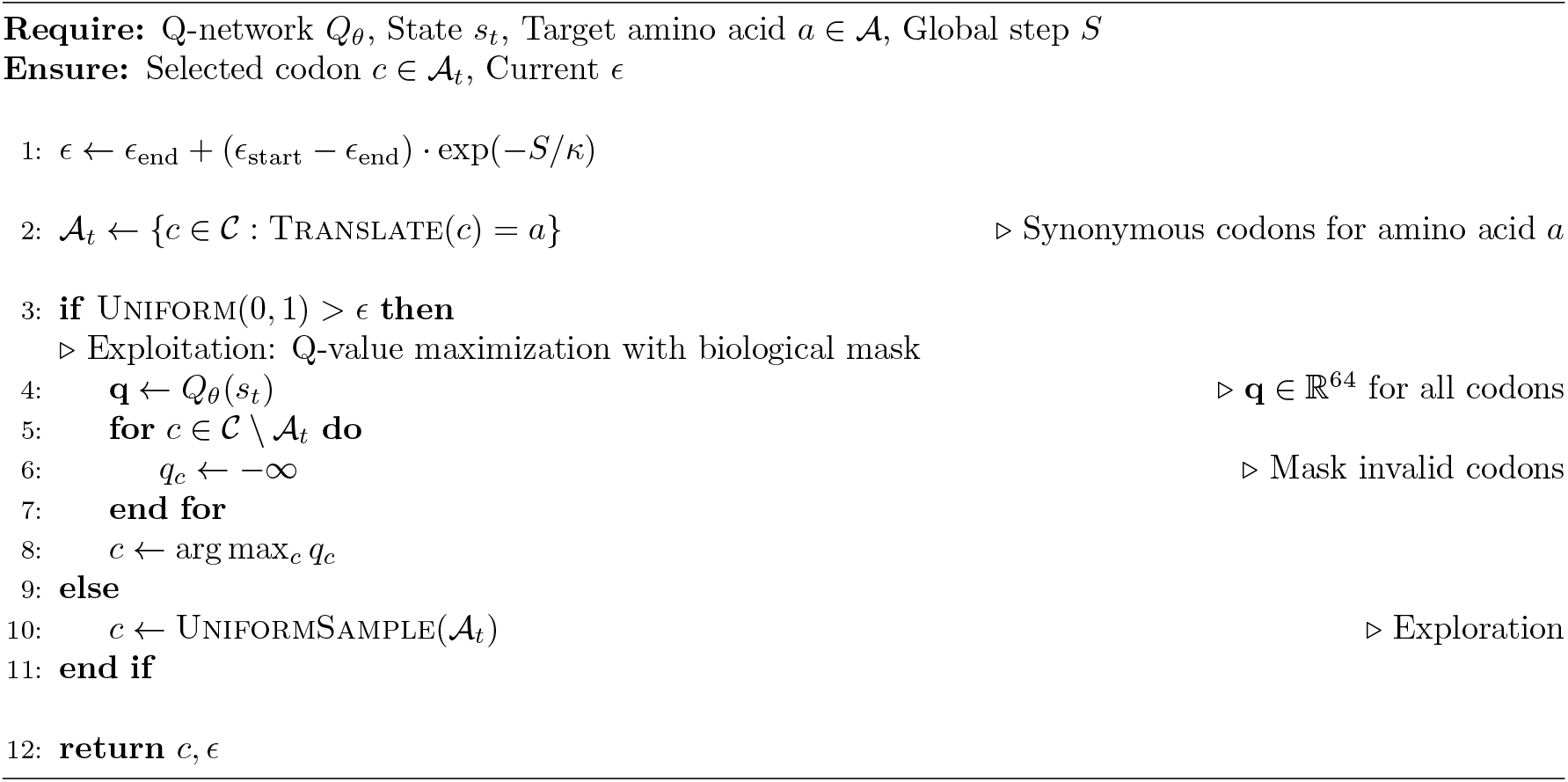

#### Algorithm 4

Bellman Optimization with Experience Replay (OptimizeBellman)

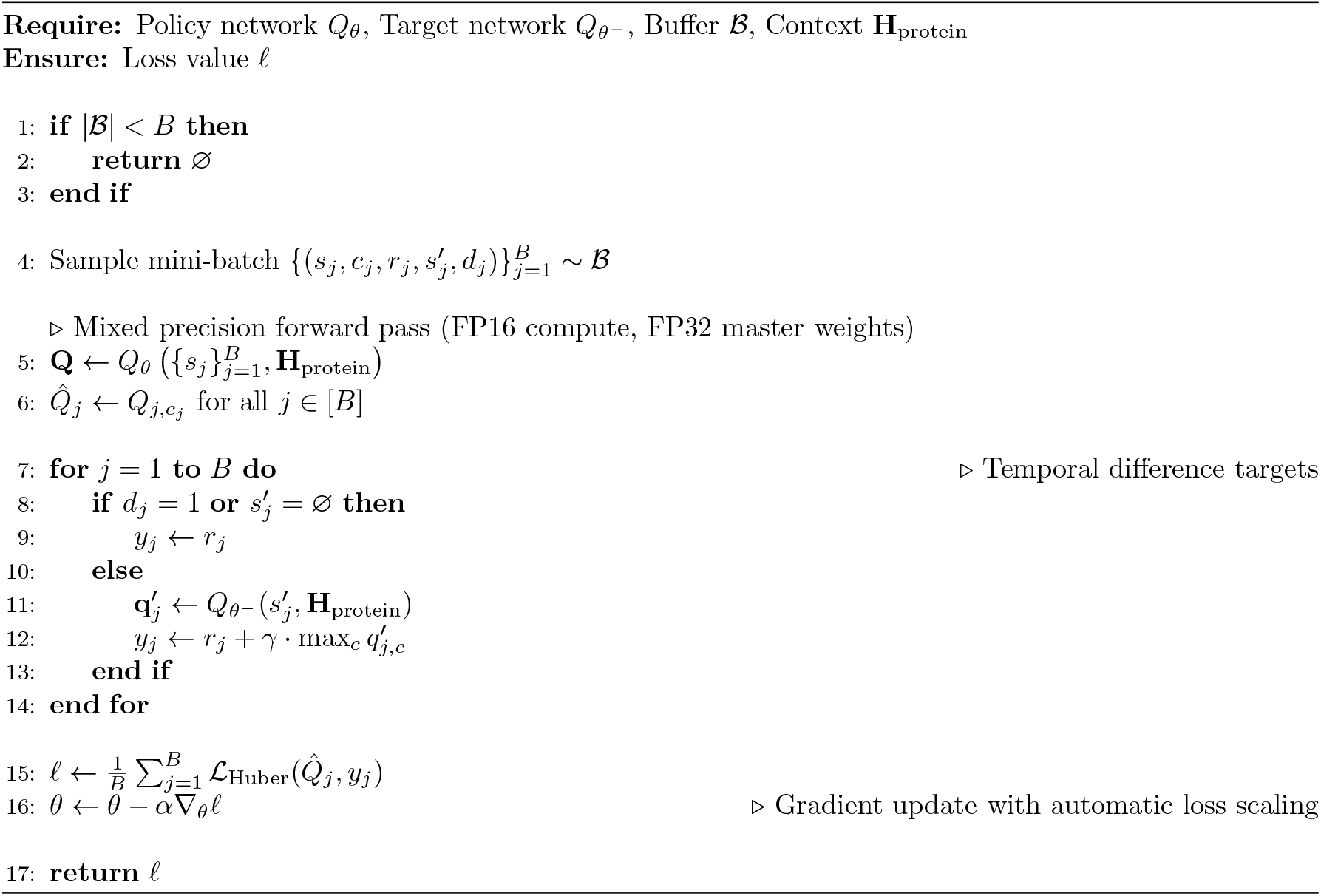

#### Algorithm 5

Multi-Objective Reward Computation (ComputeReward)

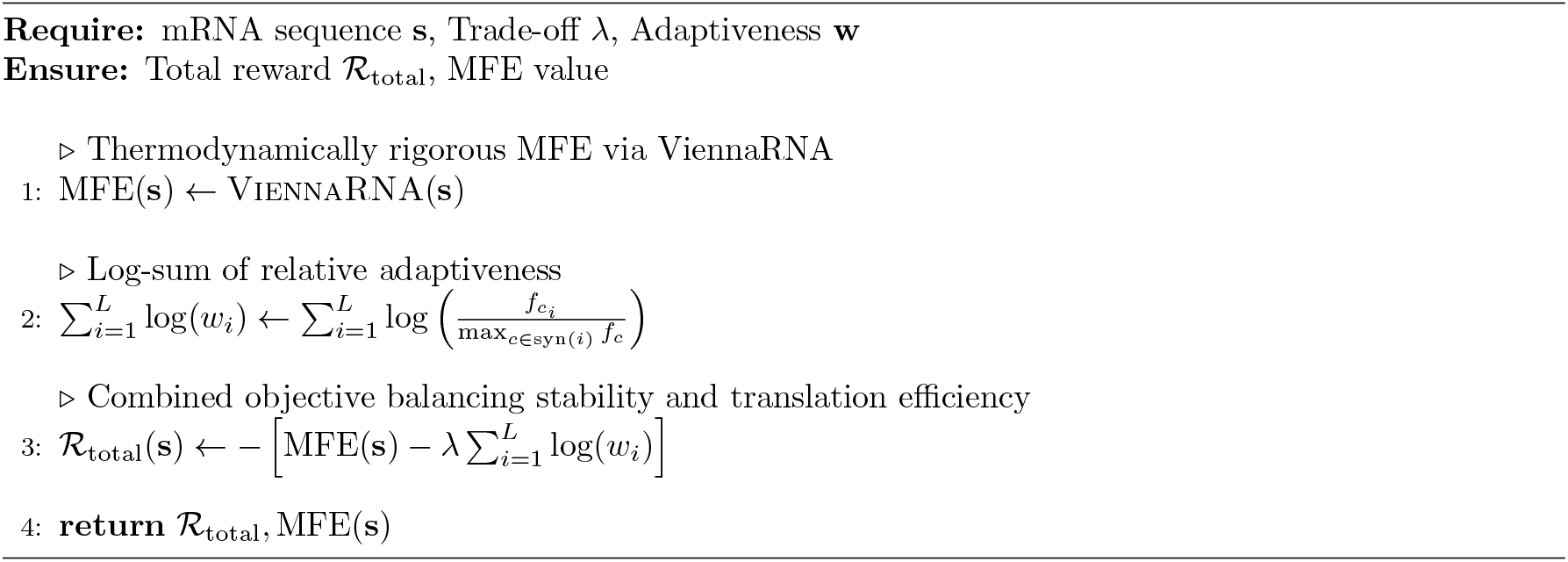

#### Algorithm 6

Milestone Reward for Reward Densification (MilestoneReward)

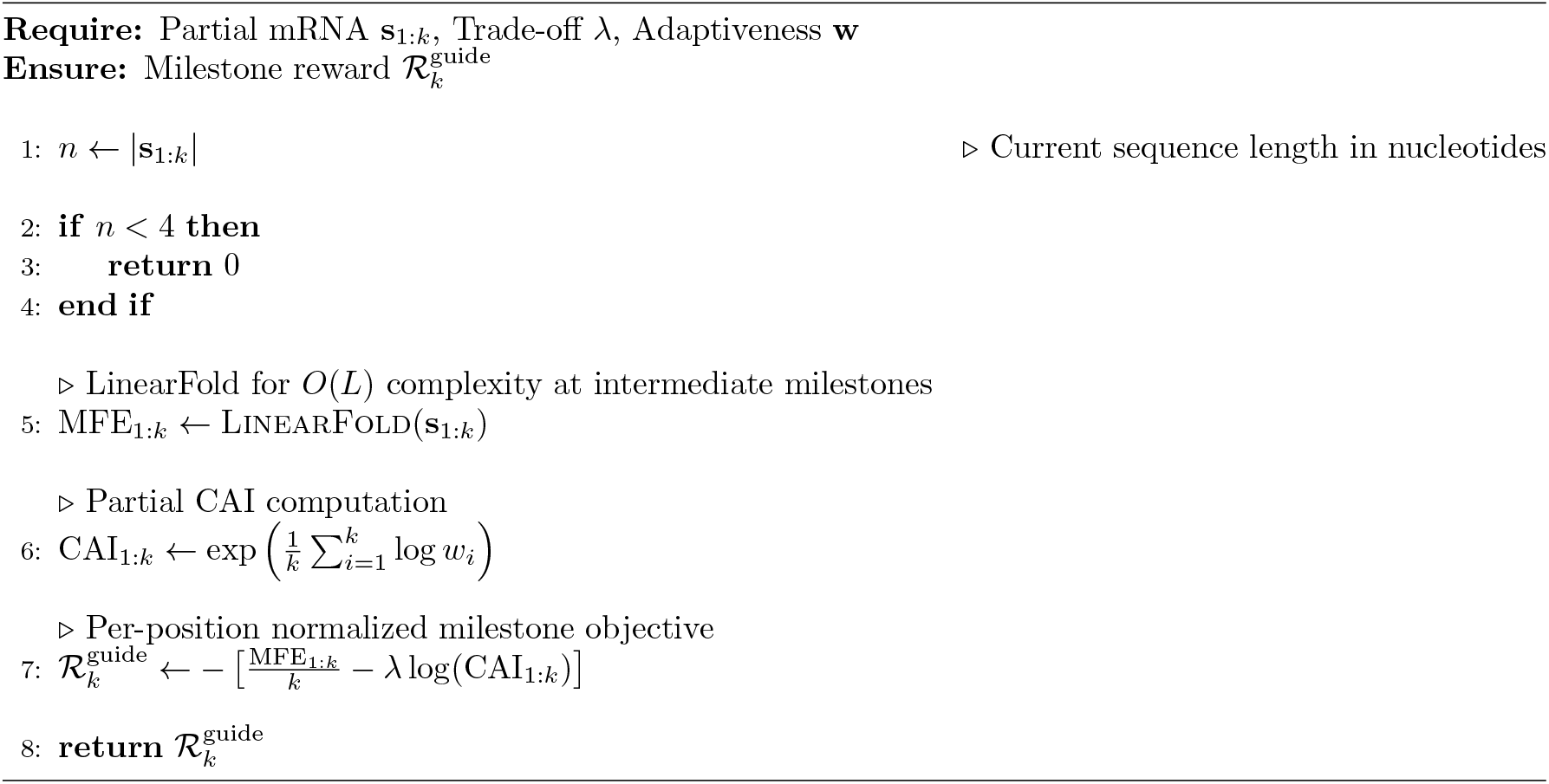

#### Algorithm 7

Replay Buffer Warm-Start with Expert Demonstrations (WarmStartBuffer)

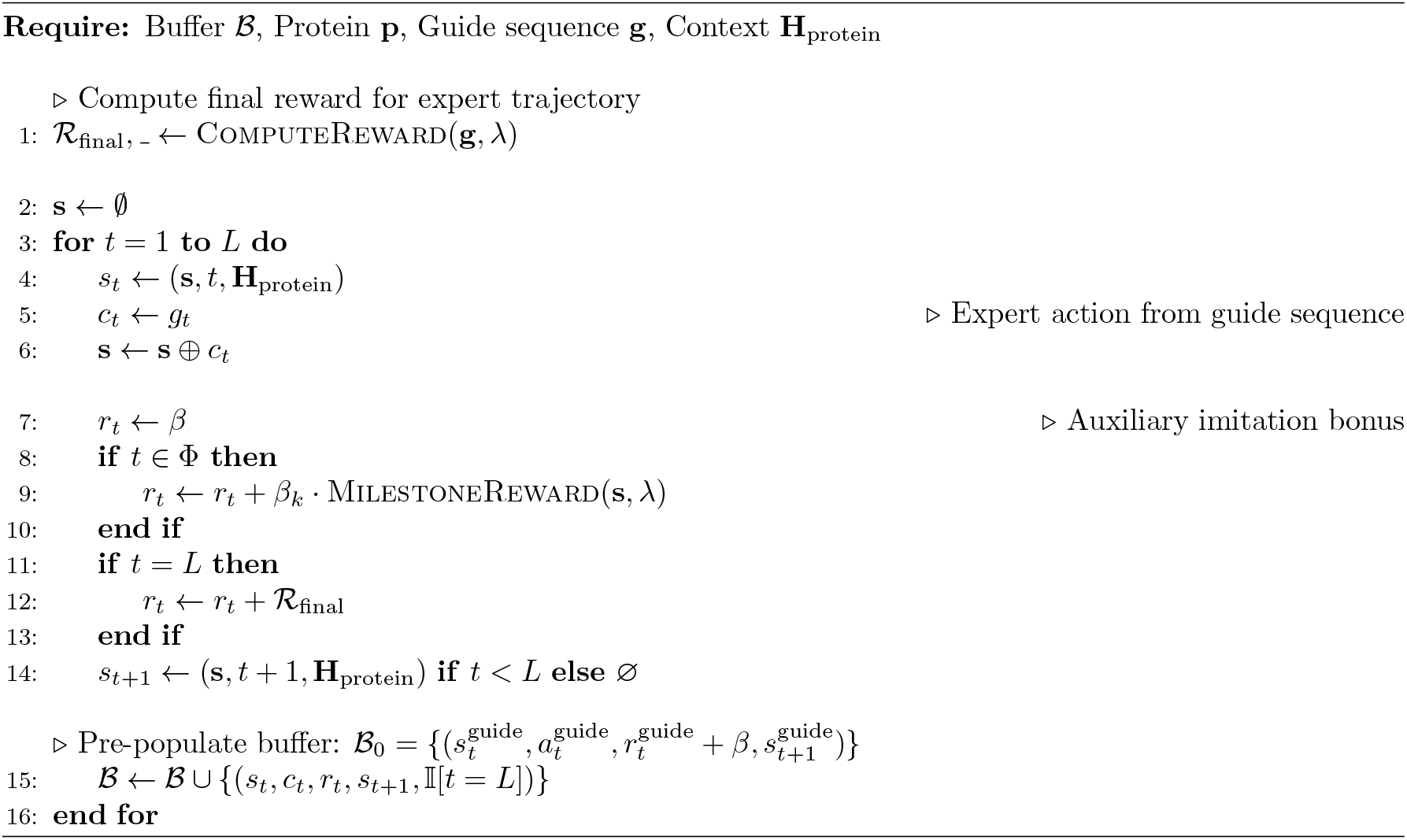

#### Algorithm 8

Flash Attention-based Q-Network Forward Pass (CodonRLForward)

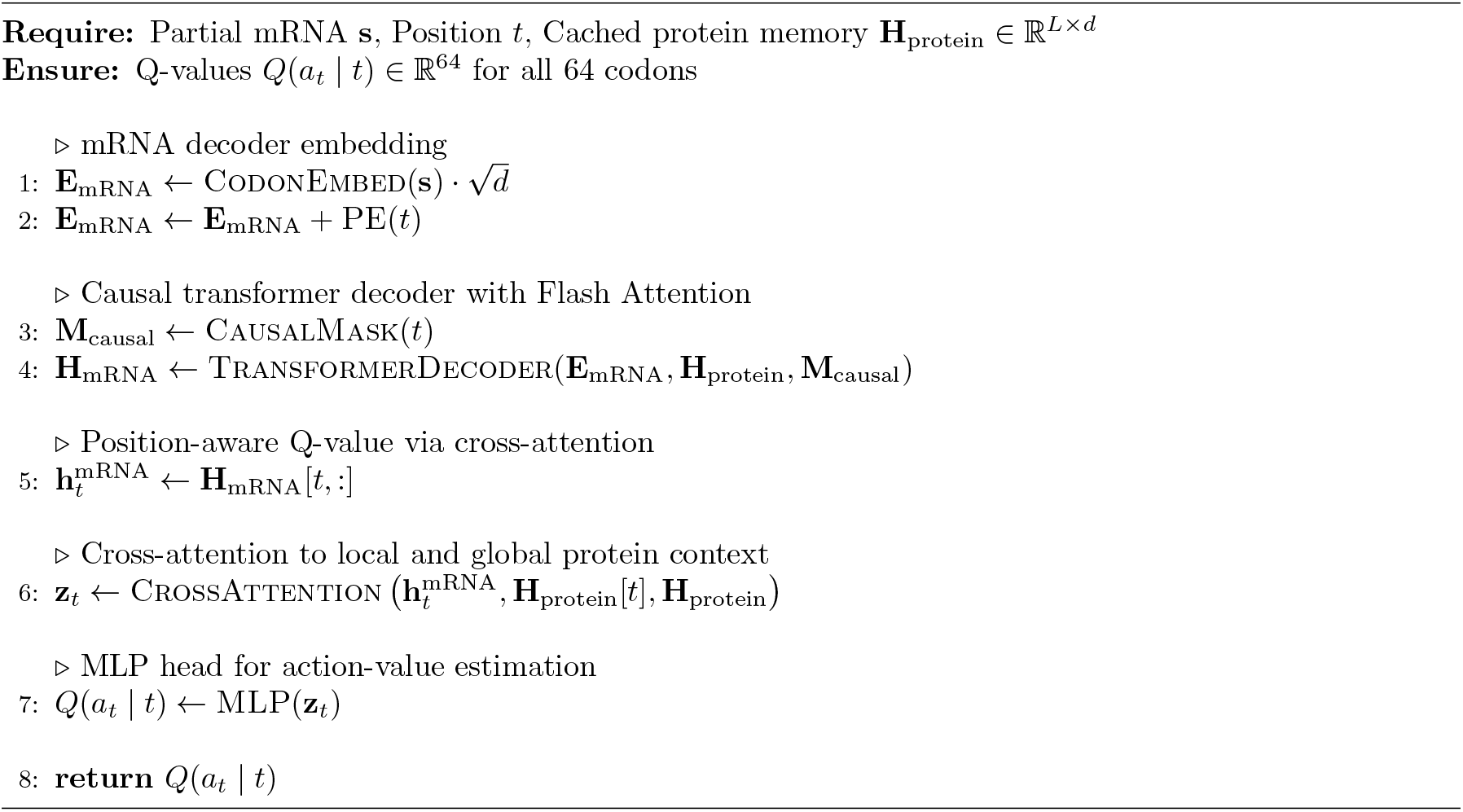

**Figure 7:**
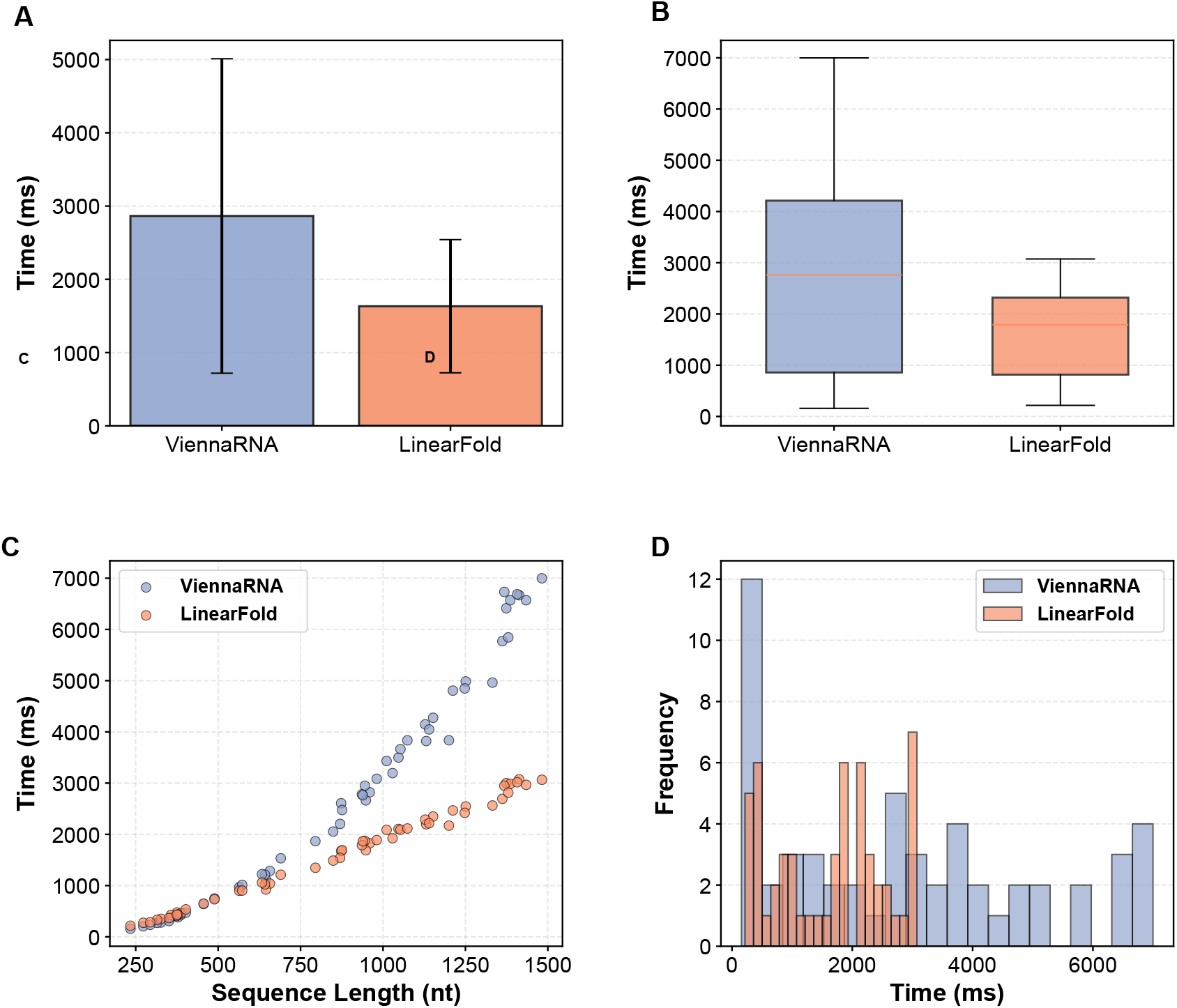
Comprehensive runtime comparison between ViennaRNA and LinearFold for MFE computation. **(A)** Mean runtime comparison. Bar plot showing the average computation time for minimum free energy (MFE) prediction using ViennaRNA (2864.90 ± 2146.16 ms, blue) and LinearFold (1633.05 ± 908.95 ms, orange). Error bars represent standard deviation (*n* = 55 sequences). **(B)** Runtime distribution. Box plots illustrating the distribution of computation times for both methods. Boxes represent the interquartile range (IQR), horizontal lines indicate medians (ViennaRNA: 2763.15 ms; LinearFold: 1789.36 ms), and red diamonds denote mean values. **(C)** Runtime vs. sequence length. Scatter plot showing the relationship between RNA sequence length (234–1482 nt) and computation time for both ViennaRNA (blue) and LinearFold (orange). Each dot represents one sequence. **(D)** Runtime frequency distribution. Overlapping histograms displaying the frequency distribution of computation times across all test sequences. LinearFold achieves 1.75*×* speedup over ViennaRNA with 57.7% lower variance (std: 908.95 ms vs. 2146.16 ms).

### Runtime Comparison between ViennaRNA and LinearFold

